# Predicting Gravitational Self-Motion: Falling Down versus Falling Up

**DOI:** 10.64898/2026.06.04.730124

**Authors:** Björn Jörges, Marlene Wessels, Heiko Hecht, Laurence R. Harris

## Abstract

Humans have been shown to use an internalized representation or prior of Earth gravity to predict object motion. Often, this is shown using the fact that humans tend to neglect an object’s acceleration when predicting it’s motion. This bias tends to be ameliorated when the acceleration acting on the object is Earth gravity. It is currently unclear whether this prior can also be used when assessing one’s self-motion. We immersed two cohorts of 20 seated participants in a virtual environment of a street scene. They experienced simulated self-motion consistent with either flying straight upwards towards the ceiling (ANTI-GRAVITY) or falling downwards towards the ground (GRAVITY). After 0.8 s to 1.4 s of motion, the screen turned blank, and participants pressed a button to indicate when they thought they reached the surface they were moving towards. In Experiment 1, they were instructed to use their eye level as the reference (i.e., when they imagined the floor or ceiling to be aligned with their eyes). In Experiment 2, they used their feet or the top of theirs heads to judge when they had reached the ceiling or floor, respectively. In Experiment 1, no differences between the GRAVITY and ANTI-GRAVITY conditions were found in terms of accuracy or precision, contrary to our hypotheses. In Experiment 2 participants pressed the button significantly later in the ANTI-GRAVITY condition than in the GRAVITY condition, in line with the predictions of the use of a gravity prior, while precision remained unaffected. Speculatively, the instructions in Experiment 1 may have made the experience less ecologically valid and immersive, thus preventing participants from activating the strong gravity prior. Experiment 2 (head/feet as reference), on the other hand, provides convincing evidence for an involvement of this prior in the prediction of self-motion. In sum, the study provides evidence that humans can rely on an internalized representation of Earth gravity when predicting self-motion, at least under ecologically valid task conditions. The absence of effects in Experiment 1 and their presence in Experiment 2 suggest that activation of this gravity prior depends on the framing of the task.

## Introduction

Humans tend to neglect the acceleration of a visual object using instead an assumption that it is moving at a constant velocity (Benguigui et al., 2003; Benguigui & Bennett, 2010; Bennett & Benguigui, 2013; Kreyenmeier et al., 2022; Wessels, Hecht, et al., 2023; Wessels, Zähme, et al., 2023; Wessels & Oberfeld, 2024) when predicting its position, likely as a consequence of the visual system’s poor sensitivity to visual acceleration (Ceccarelli et al., 2018; Werkhoven et al., 1992). When observes neglect acceleration information of a positively accelerating object, they will assume that the object reaches a predetermined point later than it is the case. Consequently, they overestimate its time-to-arrival (see also Eq. (2) in Wessels & Oberfeld, 2024). However, there is ample evidence that the perceptuo-motor apparatus is highly capable of exploiting at least one specific acceleration: that of Earth’s gravity, as revealed by our ability to relatively accurately predict the motion of falling objects (Bosco et al., 2015; Delle Monache et al., 2014, 2017; Indovina et al., 2005; Jörges et al., 2018, p. 201; Jörges & López-Moliner, 2019; Lacquaniti et al., 2015; McIntyre, Zago, & Berthoz, 2001; McIntyre, Zago, Berthoz, et al., 2001; Zago et al., 2004, 2008, 2009, 2010). This internalization of gravity has been contextualized in a Bayesian framework of perception as a strong Earth gravity prior (Jörges & López-Moliner, 2017, 2020). This prior may be acquired by humans over a lifetime of experiments in the Earth’s gravitational environment. Other evidence points towards its innateness in chicken (Bliss et al., 2023; Kobylkov & Vallortigara, 2025; Vallortigara & Regolin, 2006) even though the origin of this strong Earth gravity prior remains contested (Bardi et al., 2014; Bertenthal et al., 1985, 1987). This strong Earth gravity prior allows humans to predict object motion more accurately when an object accelerates under Earth gravity compared to when it accelerates in some arbitrary way (Zago et al., 2008).

When predicting self-motion, its acceleration component is, in contrast to the motion of external objects, and despite humans being relatively insensitive to visual self-acceleration (Schaffer & Durgin, 2005), not completely neglected: using visually simulated self-motion, Capelli et al. (2010) measured the estimated time-to-arrival for an accelerating observer to reach a predetermined point in their stationary environment after the last part of their linear forward travel was carried out in the dark. They observed judgements in between those predicted based on the observer’s final speed before occlusion (a first-order estimate) and those based on a combination of the observer’s final speed *and* their acceleration (a second-order estimate). That is, observers pressed the button earlier than would be expected under the constant speed assumption, suggesting they were at least partially sensitive to the visual consequences of their self-acceleration. Interestingly, this was only the case for accelerating self-motion; for decelerating motion, observers neglected changes in speed and predicted their motion based on the last observed speed.

This project investigates whether humans use the strong Earth gravity prior in predicting their own *falling* motion (i.e., downwards self-motion that accelerates at 1g). Relying on this prior should allow observers to compensate for the partial neglect of the acceleration component of their motion observed by Capelli et al. (2010). This should help participants make more accurate predictions of self-motion when participants accelerate in line with gravity than when their self-motion defies gravity. As in Capelli et al.’s experiment, the last part of their fall will be occluded, i.e., occur in the dark. Will they feel they continue to accelerate during this period under the influence of their strong Earth gravity prior, or will they feel that they continue to move at the last velocity they saw before the occlusion? We predict that, for gravitationally consistent simulated self-motion (falling with an acceleration of 9.8 m/s/s downwards), participants will be able to rely on their strong Earth gravity prior to predict their own falling motion correctly during occlusion as long as it is in the downward direction and consequently be relatively accurate in assessing their time of arrival at the ground. Contrariwise, when moving up, any such prior during a blanked-out phase of motion would either not be activated at all, or, if activated, tend to retard perceived acceleration and result in an overestimation of time-to-arrival.

We further expect precision (i.e., within-participant estimation variability) to be better for gravitationally consistent self-motion in comparison to self-motion under other accelerations since under Earth gravity, time-to-arrival estimates can be assisted by the strong Earth gravity prior, which is represented by the perceptual system with a higher precision (Jörges & López-Moliner, 2020) than an arbitrary acceleration (Werkhoven et al., 1992). Relying on more precise information about self-acceleration should then translate to more precise time-to-arrival judgements.

Specifically, our hypotheses are:

(1) Accuracy: Humans will overestimate their time-to-arrival when they “fall” upwards because of an incorrect impression of slowing (or maintaining constant speed) compared to when then they fall downwards, when the prior can continue to assist during the dark phase and mitigate the erroneous overestimation.
(2) Precision: Precision will be higher for gravitationally consistent falling downwards in comparison to accelerated upwards self-motion.

## Methods

### Participants

In Experiment 1, 20 participants completed the study (see power analysis in Appendix A). They were all recruited from the York University psychology undergraduate participant pool, were 18 to 24 years old, had normal or corrected-to-normal vision, and had no reported visual or vestibular disorders. All of them provided written informed consent. The study was conducted under the ethics protocol e2025-288 issued by the York University Office of Research Ethics (ORE) and followed the principles of the declaration of Helsinki. Data collection occurred in January 2026.

### Apparatus

The stimulus was programmed in Unity (v6000.0.38f1). Both the Unity project and the executable can be found in the Open Science Framework repository (https://osf.io/hyzd7/). Stimuli were presented using a PC with an Intel® Core™ i7-8700 CPU @ 3.70GHz, 16GB, 2666 MHz RAM, and an 8GB NVIDIA GeForce RTX 2070 graphics card. It was projected in an HTC VIVE Pro EYE headset (110° field of vision, 1540 × 1600 resolution per eye, and a refresh rate of 90 Hz). Participants’ responses were registered using a M-U0026 Logitech computer mouse.

### Stimulus

Participants, while seated in real life, were immersed in a virtual street environment (see insets in Figure 1) consisting of two car lanes and sidewalks lined by tall buildings. All objects were scaled to be life-sized to provide a credible and immersive environment, and the road was mirrored on the ceiling to match visual cues up and down as closely as possible. The viewpoint was simulated such that participants were facing down the street, i.e., with buildings on their right and left. During the experiment, the viewpoint of the virtual camera moved either from the ceiling downwards or from the ground upwards, simulating participants falling down or “falling up” with an acceleration of 9.8 m/s/s. After a given presentation duration, the scene was occluded and participants were tasked to press a button at the moment they thought they had reached the ground or the ceiling, respectively. In order to keep the visual stimulus equivalent in both gravity conditions, we mirrored the floor to also mark the ceiling of the scene (see Figure 1). Buildings to the right and to the left of the observer provided structural cues to scene orientation.

**Figure 1:**
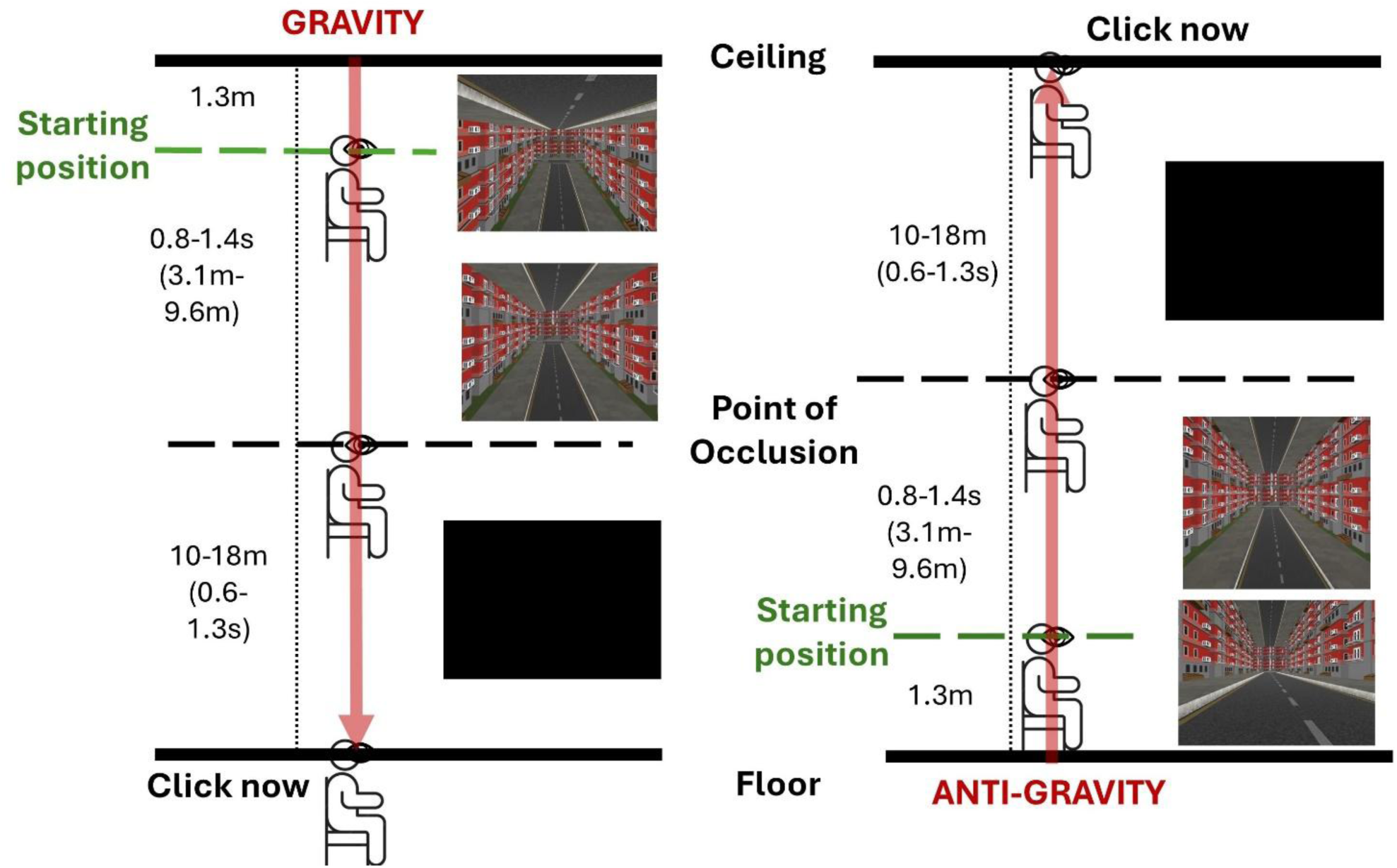
Schematic of the stimulus along with screenshots from the program for Experiment 1.

Participants experienced visual stimulation such that they either (1) started out up hovering above the scene with their eyes 1.3 m below the ceiling (see “Starting Position” in Figure 1 on the left) looking down the street and then fell down towards the road below (GRAVITY condition) with an acceleration of +9.81 m/s^2 or (2) they started out at a seated eye-height of 1.3 m above the ground (see “Starting Position” in Figure 1 on the right) and then “fell” up (ANTI-GRAVITY condition) with an acceleration of +9.81 m/s^2. After experiencing linear optic flow for one of five presentation durations (0.8 s-1.4 s in steps of 0.15 s), the screen blacked out (see “Point of Occlusion” in Figure 1) and participants had to press a button when they thought that their viewpoint had reached the target, i.e., the floor (GRAVITY condition) or the ceiling (ANTI-GRAVITY condition). Accordingly, the screen was blacked out when their viewpoint was between 10 and 18 m (in steps of 2 m) from the floor (GRAVITY) or ceiling (ANTI-GRAVITY). This is illustrated in Figure 1. The five presentation durations and five occlusion distances corresponded to occlusion durations of 0.6 s to 1.27 s. A video of the stimulus can be found on Open Science Framework (https://osf.io/hyzd7/).

### Procedure

Participants were given verbal instructions and conducted a few training trials under supervision of the experimenter until it was clear that they had understood the task. They then completed all experimental trials in one session. Specifically, the independent variables were:

- 5 presentation durations (0.8 s – 1.4 s in steps of 0.15 s)
- 2 gravity directions (GRAVITY and ANTI-GRAVITY)
- 5 occlusion distances (10 m – 18 m in steps of 2 m)

At 5 repetitions per combination of all independent variables, this made for 250 trials overall. Conditions were interleaved in a randomized fashion. Each session took no more than 20 minutes. Participants were not given any feedback about their estimation performance throughout the experimental session.

### Data Analysis

We collected temporal responses (the time during which they travelled until pressing the button) from the participants, from which we subtracted the physically correct time it would have taken the participants to reach the target. We used this signed estimation error as the dependent variable to assess differences in accuracy. The error was coded such that positive values meant that participants pressed the button too late (overestimation) and negative values meant that participants pressed the button too early (underestimation). We then excluded all trials where, across all data points, the temporal error was more than 1.5 the interquartile range below or above the first or third quartile, respectively. Using this criterion, we excluded 204 out of 5000 total trials (4%). To assess difference in precision, we computed the deviation from the mean for each combination of conditions and participant separately. That is, we subtracted the mean timing error across all five repetitions from each individual response and took the absolute value to obtain the deviation.

To assess statistical significance, we employed linear mixed modelling as implemented in the lme4 package (Bates et al., 2015) for R (R Core Team, 2017).

#### Accuracy

To test for differences in accuracy, we used the signed temporal error as the dependent variable, *gravity* (a categorical variable with the levels GRAVITY and ANTIGRAVITY), the *duration of presentation* (as continuous variable), and the *occlusion distance* (as continuous variable) as fixed effects, as well as random intercepts and random slopes for gravity, presentation duration and occlusion distance for the grouping variable Participant. The inclusion of the fixed effect “Gravity” in the model was based on our hypothesis, where we expected a difference between GRAVITY and ANTIGRAVITY, while we followed a “keep it maximal” approach for the random effects (Barr et al., 2013). The model further included the fixed effects “duration of presentation” and “occlusion distance” because, according to previously conducted simulations, representing variables as random effects but not as fixed effects can lead to inaccurate coefficient estimates. In the Wilkinson & Rogers notation (Wilkinson & Rogers, 1973), this statistical model reads as follows:

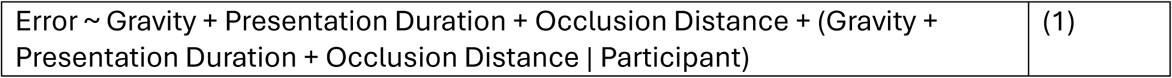

We then computed 95% confidence intervals using the confint() function from base R.

In line with our Hypothesis (1) regarding accuracy, we expected participants to press the button later in the ANTIGRAVITY condition (larger relative overestimation) than in the GRAVITY condition.

#### Precision

We used the same statistical method as for accuracy to assess differences in precision between GRAVITY and ANTI-GRAVITY, except that we used the deviation (as defined in the pre-processing section) as dependent variable.

In line with our hypothesis regarding precision, we expected the deviations to be lower in the GRAVITY condition than in the ANTI-GRAVITY condition.

You can find an analysis script as well as some pilot data in the preregistration package on Open Science Framework (https://osf.io/hyzd7/).

### Results

We found no significant differences in accuracy (b = 19 ms; 95 CI = [-22ms; 64ms]) or precision (b = 5 ms, 95 % CI = [19 ms; 8 ms]) between the gravity conditions.

The mean error (accuracy) for the GRAVITY condition was 126 ms (SD = 44 ms) and 136 ms (SD = 45 ms) for the ANTIGRAVITY condition. The mean deviations (precision) were 197 ms (SD = 171 ms) for GRAVITY and 193 ms (SD = 168 ms) for ANTIGRAVITY.

Further, higher occlusion distances were related to lower errors (at a rate of -24 ms per occluded meter, 95 % CI = [-32 ms; -15 ms]) but left precision unaffected (b = 1 ms, 95 % CI = [-1 ms; 3 ms]). Presentation duration was not related to significant differences in accuracy (b = 91 ms, 95 % CI = [-78 ms; 247 ms]) or precision (b = 1 ms, 95 % CI = [-69 ms; 51 ms]).

The results for Experiment 1 are visualized in Figure 2.

**Figure 2.**
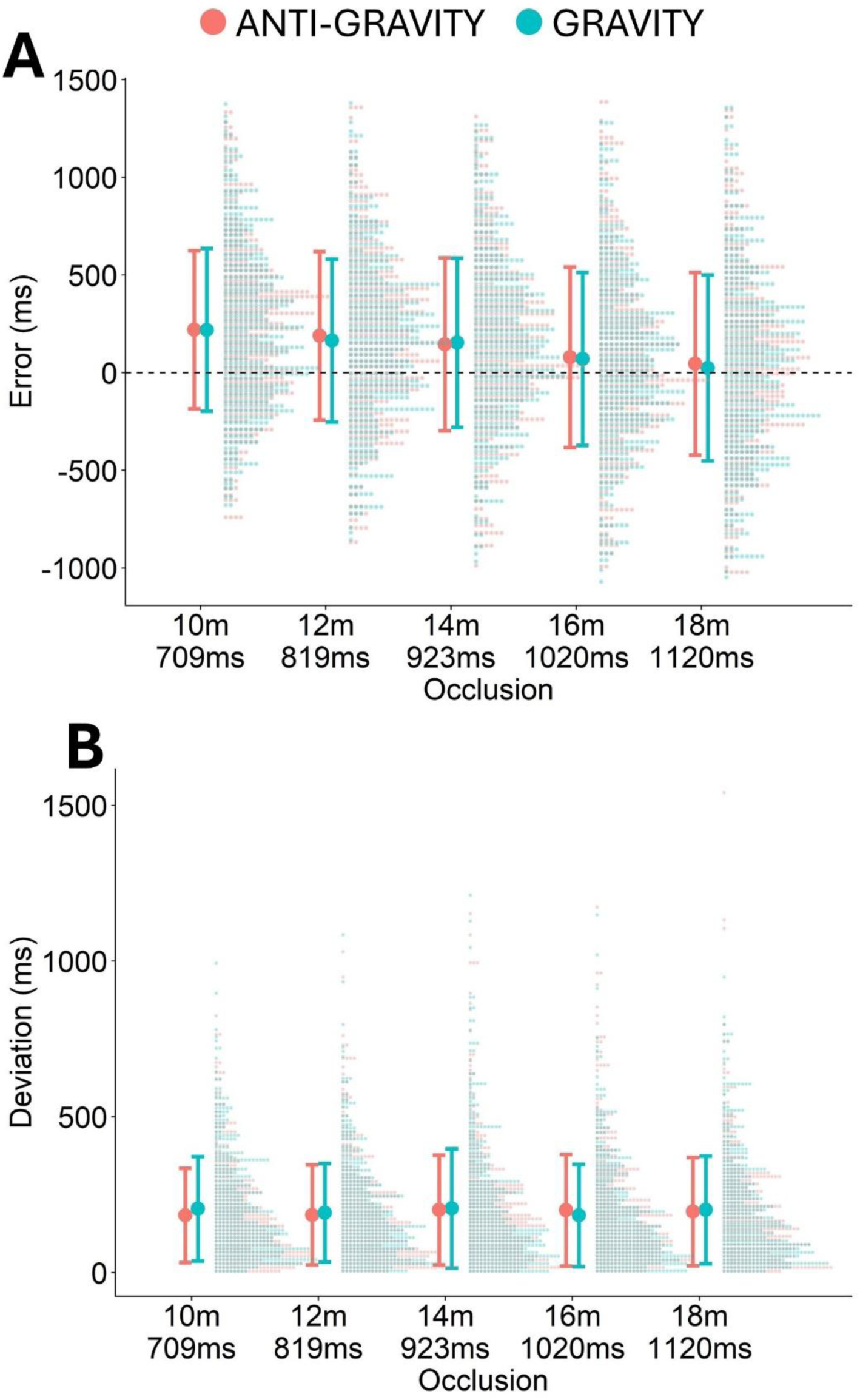
Full distributions of errors for the different occlusion distance (x axis) and the two gravity conditions (color-coded). The bold dot represents the mean across all participants and trials and the antennae indicate +- 1 standard deviation. Positive values meant that participants pressed the button too late (i.e., after they reached the target surface) and negative values meant that participants pressed the button too late (i.e., before they reached the target surface). B. Like A., but for the deviations.

## Experiment 2

In Experiment 1, we asked participants to use their eyes as reference to extrapolate their falling time, i. e., we asked them to press the button when they eyes crossed the reference surface. This may have been unintuitive and may have interfered with their sense of realism and immersion. especially for the falling condition where the right answer would have involved them being heavily immersed in the tarmac. We therefore repeated the Experiment but asked participants to use their feet (for GRAVITY) and the top of their head (ANTI-GRAVITY) as references. We collected a new set of 20 participants, as per the power analysis conducted for Experiment 1.

### Participants

A new set of 20 participants was recruited from the York University Undergraduate Participant Pool. They were 18 to 31 years old. Their remaining characteristics matched the description of the cohort tested for Experiment 1. Data collection for this experiment occurred in February and March 2026.

### Stimulus and Procedure

We changed the scene and the motion parameters such that the distance participants were falling remained matched between both conditions (see Figure 3). In the ANTI-GRAVITY condition, the viewpoint started 1.3 m above the floor (average seated eye-height) and the ceiling was simulated at a height of 1.3 m + visible distance + occluded distance + 0.1 m (distance between eyes and top of the head). In the GRAVITY condition, the viewpoint started 1.3 m below the ceiling, which in turn was simulated at a height of 1.3 m (buffer between head and ceiling at motion onset; added such that the distance between eyes and starting surface was matched between GRAVITY and ANTIGRAVITY)) + visible distance + occluded distance + 1.3 m (distance between feet and eyes while seated). No other parameters were changed.

**Figure 3.**
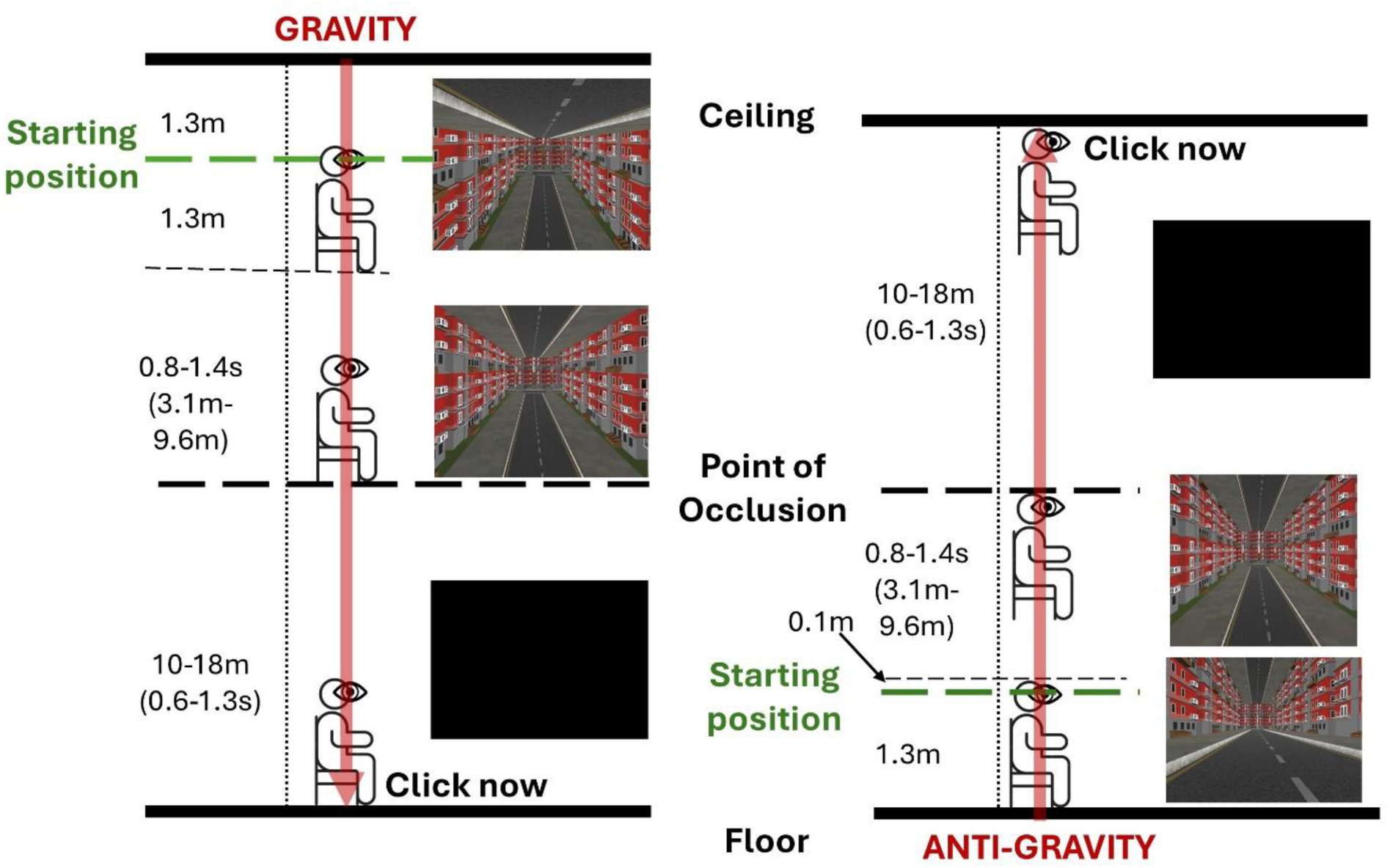
Schematic of the stimulus including screenshots from the program, updated for Experiment 2. Please note that, as indicated by the diagram, the room height was not matched between GRAVITY and ANTIGRAVITY (in favor of matching the distance between the viewpoint and the closer surface before motion onset), but both the falling distance and the occluded distance were matched. See the Discussion section for an examination of possible introduced by this decision.

### Results

Participants pressed the mouse button significantly later for ANTI-GRAVITY than for GRAVITY (by 65 ms, 95 % CI = [29 ms; 102 ms]). This is visualized in Figure 4A. Deviations were numerically higher for ANTI-GRAVITY than for GRAVITY (by 8 ms), but not significantly so (95% CI = [-7 ms; 20 ms]) – see Figure 4B.

**Figure 4.**
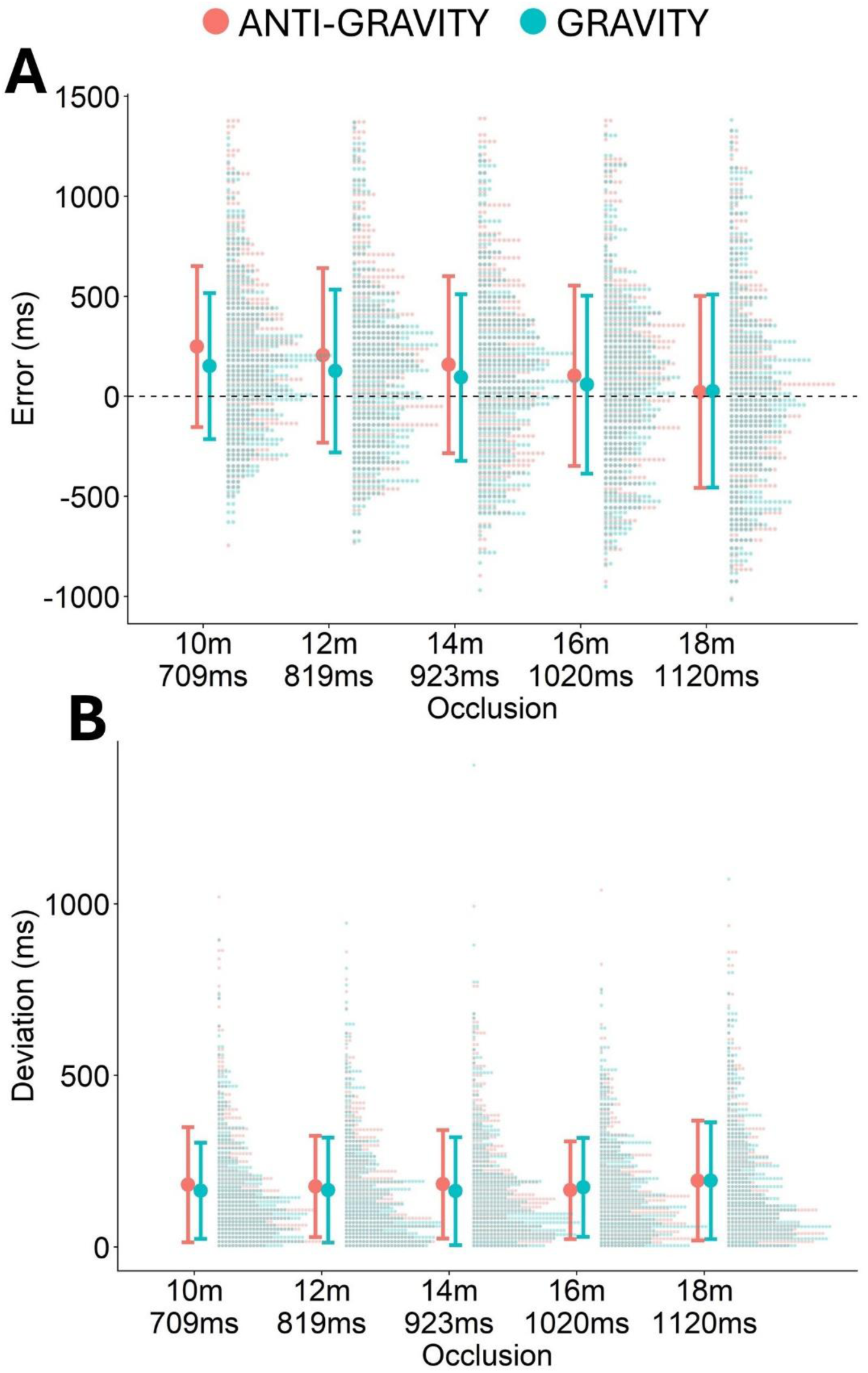
A. Full distributions of errors for the different occlusion distance (x axis) and the two gravity conditions (color-coded). The bold dot represents the mean across all participants and trials and the antennae indicate +- 1 standard deviation. B. Like A., but for the deviations.

The mean error (accuracy) for the GRAVITY condition was 92 ms (SD = 43 ms) and 148 ms (SD = 45 ms) for the ANTIGRAVITY condition. The mean deviations (precision) were 172 ms (SD = 154 ms) for GRAVITY and 180 ms (SD = 159 ms) for ANTIGRAVITY.

Each meter of occluded distance decreased the error by 21ms (95 % CI = [13 ms; 30 ms]), but precision was not significantly related to occluded distance (b = 2 ms, 95% = [-0.2 ms; 4 ms]). No significant relationship was found between presentation duration and accuracy (b = -95 ms, 95 % = [-206 ms; 18 ms]) or precision (b = -6 ms, 95 % = [-43 ms; 27 ms]).

## Discussion

In this project, we tested whether humans used the strong gravity prior in the prediction of self-motion. Such a prior has been previously found to be recruited in predicting the motion of falling objects (e.g., McIntyre, Zago, Berthoz, et al., 2001; Zago et al., 2008) the human perceptual apparatus tends to neglect arbitrary accelerations and rely on first-order (i.e., speed-based) information when predicting object motion in the visual domain. When an object accelerates, however, Earth gravity has a special status, and humans can rely on knowledge of this specific acceleration to predict an object’s motion more successfully.

Accordingly, we had expected participants to judge that they had reached the target surface earlier when their fall was consistent with gravity than when they fell “up” as in anti-gravity, in line with the idea that the gravity prior could be used to compensate for the partial neglect of acceleration information. We found no evidence for this hypothesis in our first experiment where participants were asked to predict their physical motion using their eyes as the reference. In a second experiment, where participants used the soles of their feet and the top of their head as reference points, the mouse button was pressed significantly later for ANTI-GRAVITY than for GRAVITY (by 65 ms), in line with our strong gravity prior hypothesis. This suggests that the strong gravity prior is activated not only for the prediction of object motion, but also for the prediction of self-motion, aiding us, for example, to better time our landings after jumping.

To our knowledge, there is only one other study that has investigated the strong gravity prior or internal model of gravity for self-motion in a different domain (Delle Monache et al., 2025). The authors found motion intervals to be judged as shorter when observers experienced (physical) downwards self-motion than for upwards self-motion in absence of visual input. This is complimentary with our project as it supports the same underlying idea, i. e., the fact that our knowledge of gravity is used in estimating self-motion, with a vastly different methodology that included a different performance measure (duration judgements versus motion extrapolation), a different input modality (vestibular versus visual), and a different acceleration profile (a sinusoidal acceleration pattern versus constant accelerations at 1 g/-1 g). Like in the present study, Delle Monache et al. (2025) did also not find any differences in precision.

The accuracy difference between GRAVITY and ANTIGRAVITY shows that the manipulation worked in Experiment 2. This makes it all the more surprising that we did not find the expected difference in precision. The gravity prior should, if recruited, be available as an additional source of information about the motion dynamics, and the presence of additional information should lead to an increase in precision. One potential explanation is that other sources of variability dominated: under the assumptions of our power analysis (see Appendix A), we would expect a mean deviation of 67 ms for GRAVITY and 85 ms for ANTIGRAVITY, versus observed values of 172 ms and 180 ms in Experiment 2. This much-higher-than-expected variability may have obscured the positive impact of recruiting the strong Earth gravity prior.

The main difference between Experiment 1 and Experiment 2 was the reference participants used to judge their arrival at the target surface. In Experiment 1, participants were asked to press the mouse button when their eyes reached the reference surface. This may have been unintuitive, in particular for the GRAVITY condition, where they would have had to imagine themselves as falling part way through the ground, thus breaking immersion. In line with this notion, previous studies have shown that the recruitment of the gravity prior may be dependent on the visual context like a 2D scene that indicates where “up” is with pictorial cues (Zago et al., 2011). In Experiment 2, we therefore instructed participants to use their feet and the top of their head as references and adjusted the stimulus such that the presentation duration (before occlusion) and the occluded distance were matched between GRAVITY and ANTI-GRAVITY. Numerically, performance in the ANTI-GRAVITY condition remained stable between Experiment 1 (with an average error of 136 ms) and Experiment 2 (148 ms). Rather, the difference between the gravity conditions in Experiment 2 emerged because the average error was much lower for GRAVITY in Experiment 2 (92 ms) than in Experiment 1 (126 ms). Interestingly, the errors decreased with increased occlusion durations; this is in line with the idea that our visuomotor TTC system may be calibrated to an anticipation time of about 1 second (see, e.g., (Hecht et al., 2002; Oberfeld et al., 2014)).

The choice to use head and feet as reference induced several potential minor confounds:

- While the falling distance was matched between the conditions, the distance between floor and ceiling was 120 cm (or around 5 % for the average trial) larger for GRAVITY than for ANTI-GRAVITY trials. This might have a hard-to-predict influence on predicted falling durations if participants use the perceived overall height of the scene to interpret their self-motion. However, a 5% difference is well below threshold and should therefore only have a minor impact.
- The point of occlusion was also not fully matched. For GRAVITY, it was at the occluded distance + 130 cm above the floor, while for ANTIGRAVITY, it was at the occluded distance + 10 cm below the ceiling. Effects that bias perceived distance, such as the well-documented compression of space in VR (El Jamiy & Marsh, 2019), might then affect the two conditions differently. However, at the average compression effect size of 10 %, this would only add 12 cm to the perceived occluded distance. This in turn corresponds to only 5 to 7 ms in added predicted occluded duration when using the values from the power analysis to model this effect – much lower than the mean difference observed for Experiment 2 (65 ms).
- Most obviously, and by design, the imagined collision point on the body was not matched between GRAVITY (feet) and ANTI-GRAVITY (top of the head). It seems possible that the imagined collision point may have shifted either towards the center of the body or towards the viewpoint for some participants. Since the correct reference for GRAVITY is further away from both the center of the body and the viewpoint, the perceived occluded distance in this condition would have, if anything, been increased more for GRAVITY than for ANTI-GRAVITY. This would increase the predicted falling duration, which is opposite to the strong gravity prior predictions and our results. The opposite bias, i.e., shifting the imagined collision point outside of the body (below the feet for GRAVITY or above the head for ANTI-GRAVITY), seems *a priori* much less likely.

In sum, we found evidence for a specific prior that relates to earth gravity, only when physically possible and believable motion was simulated. While the explanation that the counter-intuitive collision reference in Experiment 1 broke participants’ immersion, thus disabling their access to the strong gravity prior, is appealing, we did not survey participants’ immersion or presence in either of the two experiments. More research into the relationship between a strong gravity prior-induced bias and immersion or presence is thus required. This may be supplemented well by increasing immersion even further with concurrent vestibular stimulus, e.g., using motion platforms.

## Open Science

All data, materials, and scripts as well as the preregistrations can be found on Open Science Framework (https://osf.io/hyzd7/).

We deviated in the following ways from the preregistration:

- We fixed an error in the preregistered analysis script (for both experiments) in order to accurately match the outlier analysis as described in the preregistration manuscript.
- We did not conclude data collection for two participants enrolled for Experiment 2; the first consistently pressed the button so late that they were only half-way through the experiment when their timeslot was over, and the second kept the mouse button pressed continuously rather than reacting to the stimulus despite having the instructions explained to them multiple times.
- We corrected wording in the methods suggesting that participants were facing one of the walls while experiencing self-motion, to match the actual task during which participant were facing down the street.

## Appendix A Power Analysis

We used a power analysis by simulation approach (see e.g., (Debruine & Barr, 2021; Jörges, 2021). In order to simulate participant response, we assumed that participants would estimate the time it took from the point of screen blanking to the target based on:

- The remaining distance

- Assumed to be estimated accurately in both Gravity conditions with a Weber Fraction of 5%
- Their speed at disappearance

- Assume to be estimate accurately in both Gravity conditions with a Weber Fraction of 5%
- Their acceleration

- Assumed to be estimated accurately at Earth gravity and a Weber Fraction of 10% (Jörges & López-Moliner, 2020) in the GRAVITY condition
- Assumed to be estimated at 70% of Earth gravity and a Weber Fraction of 20% (Jörges et al., 2018) in the ANTI-GRAVITY condition

For each of these variables, we erred on the side of caution by choosing conservative values in the face of uncertainty, i.e., higher values in the case of Weber Fractions to simulate slightly higher-than-expected variability and a lower-than-expected accuracy difference between Gravity and Anti-Gravity.

We then simulated 100 datasets for a range of repetitions per condition and number of participants and analyzed each of these simulated data sets using the process outlined in the Data Analysis section. We recorded the p values for the difference between GRAVITY and ANTI-GRAVITY (approximated using the lmerTest package; Kuznetsova et al., 2017) for each iteration and used the fraction of significant p values (at p < 0.05) as a proxy for the expected power for any given combination of repetitions and number of participants. Even for the worst case we simulated (10 participants with two repetitions each), the expected power was > 0.99 both for accuracy and precision. As a matter of precaution, we still opted for a larger sample size of 20 participants and four repetitions each.

The R script used for the power analysis can be found on Open Science Foundation (https://osf.io/hyzd7/).

## Notes

### Competing Interest Statement

The authors have declared no competing interest.

